# Structural basis of HMCES interactions with DNA reveals multivalent substrate recognition

**DOI:** 10.1101/519595

**Authors:** Levon Halabelian, Mani Ravichandran, Yanjun Li, Hong Zheng, L. Aravind, Anjana Rao, Cheryl H Arrowsmith

## Abstract

HMCES can covalently crosslink to abasic sites in single-stranded DNA at stalled replication forks to prevent genome instability. Here, we report crystal structures of the HMCES SRAP domain in complex with DNA-damage substrates, revealing interactions with both single-stranded and duplex segments of 3’ overhang DNA. HMCES may also bind gapped DNA and 5’ overhang structures to align single stranded abasic sites for crosslinking to the conserved Cys2 of its catalytic triad.

---

DNA bases are constantly damaged by factors such as reactive oxygen species (ROS), chemotoxic agents, ionizing radiation (IR), and UV radiation^1^, and are subject to physiological modification by enzymes such as AID, the TETs and DNA methylases^2,3^. These types of DNA alterations are primarily repaired by the base excision repair (BER) pathway, which is initiated by DNA glycosylases that recognize and cleave damaged or modified bases creating apurinic/apyrimidinic sites (AP sites) or abasic sites^1^. HMCES was recently reported to recognize and covalently crosslink to abasic sites in single-stranded DNA (ssDNA), generated by uracil DNA glycosylase (UDG) at stalled replication forks^4^. The authors suggested that these DNA-protein crosslink (DPC) intermediates prevented ssDNA breaks that may consequently occur upon cleavage by AP endonucleases, which could subsequently be repaired through error-prone pathways4.

Human HMCES has a highly conserved N-terminal SOS Response-Associated Peptidase domain (SRAPd) that is widely found in bacteria and eukaryotes, with occasional presence in certain bacteriophages and archaea^5^. Animal SRAP proteins have an additional C-terminal disordered extension with multiple copies of the PCNA-interacting motif (PIP)^4^. Gene-neighborhood analysis identified SRAPd as a novel component of the bacterial SOS response, associated with multiple components of the DNA repair machinery^5^. The SRAPd contains a highly conserved triad of predicted catalytic residues, namely Cys2, Glu127, and His210, which are believed to support autoproteolytic activity^5^. In addition, Cys2 was recently shown to mediate the DPC activity^4^. To better understand the mechanism of HMCES association with DNA, we crystallized the human HMCES SRAPd in its DNA-free form (Apo-SRAPd) and in complex with several DNA-damage substrates containing 3’ overhangs of different lengths.

The crystal structure of SRAPd in complex with duplex DNA containing a three-nucleotide overhang at the 3’ end (referred to here as SRAPd_3nt) revealed SRAPd binding to two DNA molecules: DNA-A interacts via the 3’ overhang, and another molecule (DNA-B) via the blunt-end (Fig. 1a). Both DNA interaction surfaces are highly conserved.

**Figure 1.**
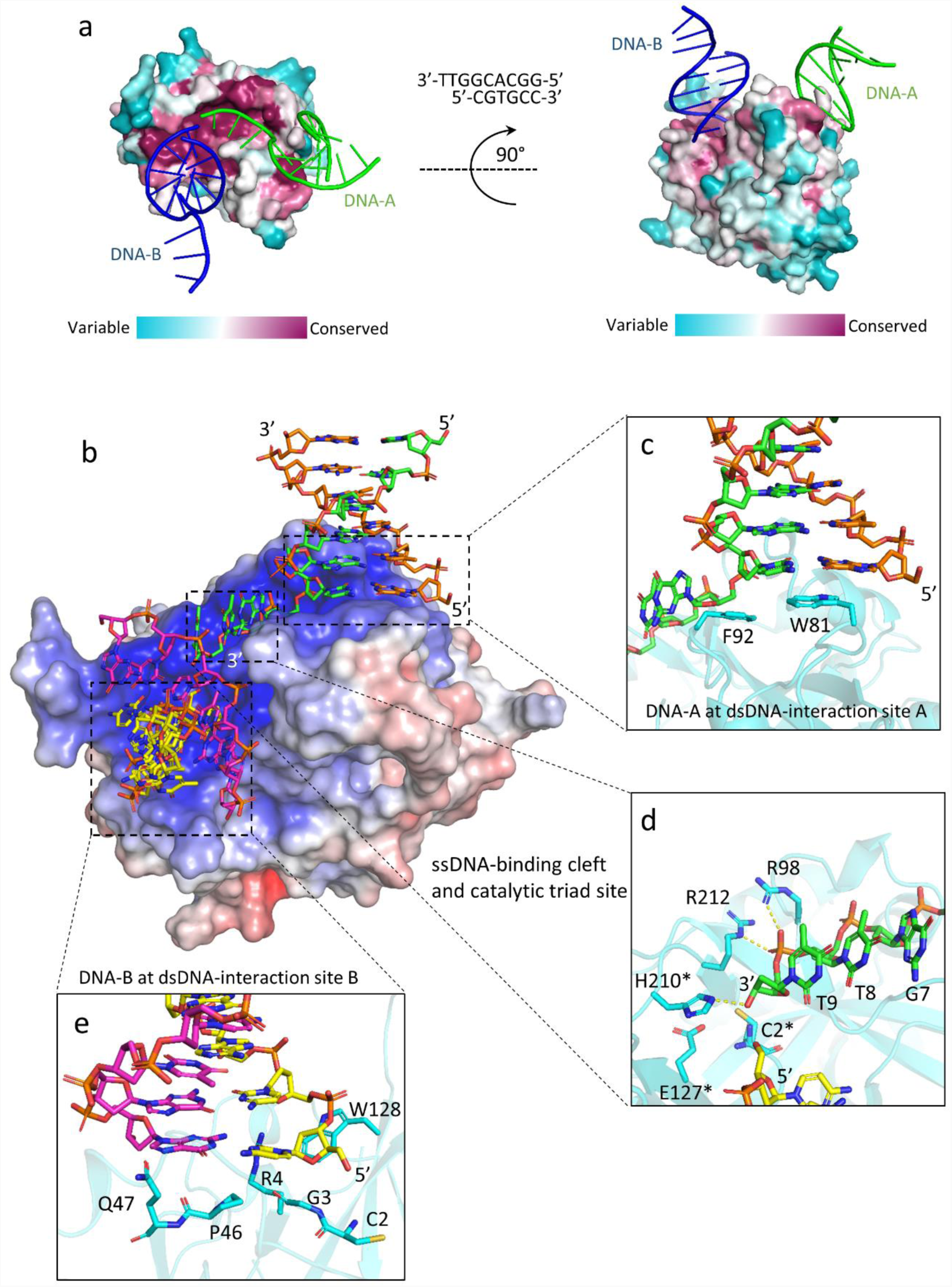
Interactions between SRAPd and 3’overhang DNA. **(a)** Surface representation of SRAPd colored by degree of sequence conservation, bound to two symmetry-related DNA molecules shown in green (DNA-A) and blue (DNA-B). Evolutionary conservation was assessed by ConSurf web server^9^. The 3’ overhang DNA sequence that was used in co-crystallization is shown in the middle. (**b)** Electrostatic surface potential representation of SRAPd interacting with two symmetry-related DNA molecules: DNA-A in green and orange, and DNA-B in magenta and yellow. SRAPd surface color indicates electrostatic potential ranging from −7kT/e (red) to +7kT/e (blue). Electrostatic surface potentials were calculated using APBS^10^. (**c)** Close-up view of the SRAPd dsDNA-interaction site A in stacked conformation with the duplex segment of DNA-A. (**d)** Close-up view of the SRAPd catalytic triad site and ssDNA-binding cleft bound to the phosphate backbone of single-strand segment of DNA-A. The catalytic triad residues as well as R98 and R212 in the ssDNA-binding cleft are shown as stick models in cyan. The ssDNA segment of DNA-A is shown in stick model in green. The DNA-B molecule is shown in stick model in yellow. The catalytic triad residues are marked with asterisks. (**e)** Close-up view of the SRAPd dsDNA-interaction site B, which stacks with the blunt-end of DNA-B.

SRAPd interacts with the 3’ overhang of DNA-A through a hydrophobic shelf created by Trp81 and Phe92, which form pI stacking interactions with the duplex segment of DNA at the ssDNA-dsDNA junction (**Fig. 1b, c**). The ssDNA 3’ overhang is sharply bent by ∼90 degrees and lies in a narrow, positively charged cleft directing it towards the catalytic triad. The ssDNA-binding cleft includes conserved Arg98 and Arg212, which form salt-bridges with the phosphate backbone of ssDNA (**Fig. 1b, d**). Alanine substitutions of either of these Arg residues severely hinder ssDNA-binding (**Supplementary Fig. 1**), and are consistent with gel-shift assays reported by Mohni et al.^4^

The pocket housing the catalytic triad accommodates the 3’-OH of the ssDNA overhang (**Fig. 1d**). Mutating the catalytic triad residues independently yielded SRAPd variants with higher affinity for ssDNA compared to wild type (WT) protein, suggesting a role other than simply DNA binding (**Supplementary Fig. 1**). In the SRAPd_3nt structure, Cys2 is ∼5.0Å from Thymine9 (T9) of the 3’ overhang (**Fig. 2a**). In the case of an abasic site, the deoxyribose moiety could freely rotate to within less than 3Å to crosslink with Cys2 at the catalytic triad site (**Fig. 2a**). This observation provides the structural logic for the recent demonstration of HMCES sensing and covalently crosslinking to abasic sites in ssDNA through Cys2^4^.

**Figure 2.**
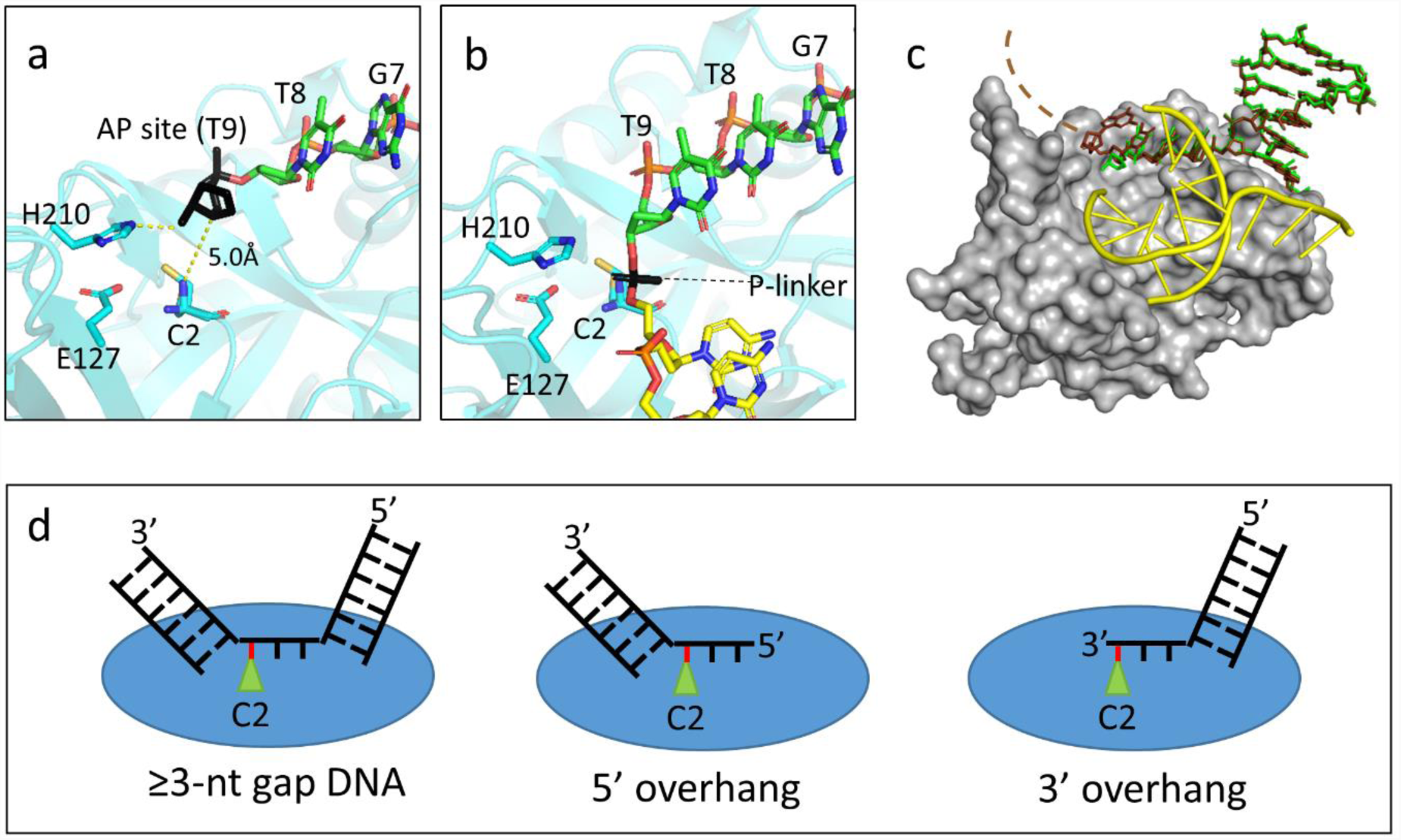
Potential DNA damage/repair substrates for SRAPd. **(a)** Close-up view of the catalytic triad site, where Thymine9 of the 3’overhang is replaced with an abasic site for illustration purposes and colored in black. The distance between the abasic site and Cys2 is labeled in Angstroms. (**b)** Close-up view of the SRAPd catalytic triad site indicating that the 3’ and 5’ ends of the two DNA molecules could be linked by a phosphate group, shown in black color for the purposes of illustration. (**c)** Crystal structure of SRAPd in complex with duplex DNA containing a six-nucleotide 3’ overhang. SRAPd is shown in surface representation in grey. The six-nucleotide overhang at the dsDNA-interaction site A is shown in stick model in brown superposed with SRAPd_3nt three-nucleotide overhang, in green. The electron density for the last two nucleotides in the six-nucleotide overhang was not resolved and hence it is not modeled (indicated by dashed lines). DNA-B at the dsDNA-interaction site B is shown in cartoon representation in yellow. (**d)** A model illustrating the potential DNA damage substrates that can be recognized by HMCES. Red line represents the abasic site in DNA.

Mohni et al.^4^ showed that HMCES forms DPC intermediates with abasic sites in ssDNA generated by uracil-DNA glycosylase (UDG), which is a monofunctional glycosylase that cannot cleave ssDNA. However, other variants of damaged bases require the use of bifunctional glycosylases with both glycosylase and lyase activities, such as NEIL3, which is a single-strand specific glycosylase with a limited lyase activity able to cleave ssDNA 3’ to an abasic site to generate a 3’ overhang^6^. Our structure indicates how SRAPd can recognize and potentially crosslink to abasic sites at the 3’ end of ssDNA overhangs (**Fig. 2a, d**). By forming a stable DPC, SRAPd is thought to protect the ssDNA from error-prone DNA synthesis and nucleolytic degradation, thus safeguarding genome integrity^4^.

Our SRAPd_3nt structure also revealed that the blunt-end of DNA-B interacts with SRAPd via dsDNA-interaction site B, composed of residues Gly3, Arg4, Pro46, Asp47, W128 (**Fig. 1e**). This interaction surface represents a potential binding site for 5’ overhang DNA, as SRAPd was shown to bind both 5’ and 3’ overhangs with similar affinities^4^. This dsDNA-interaction site B accounts for the remaining highly conserved residues of the SRAPd, suggesting that it is a universal functional feature of this domain. It is immediately adjacent to the catalytic triad and forms a contiguous, similarly charged surface with the ssDNA binding site (**Fig. 1b**). These features suggested that dsDNA-interaction site B may also be able to accommodate ssDNA extending from a longer 3’ overhang substrate bound to the dsDNA-interaction site A.

To address this question, we determined the crystal structure of SRAPd with DNA containing a six-nucleotide overhang at the 3’ end (referred to here as SRAPd_6nt). Although SRAPd has 10-fold higher affinity for ssDNA compared to dsDNA (**Supplementary Fig. 1**), the longer 3’ overhang did not displace the blunt-end-interacting DNA-B from its dsDNA-interaction site B. Instead, the extra single strand bases protrude out of the catalytic triad pocket (**Fig. 2c**), (**Supplementary Fig. 2**). This suggests that the dsDNA-interaction site B has been specifically evolved to bind duplex DNA and may form the binding site for 5’ overhang DNA structures as well. Nevertheless, given that DNA is a mediator of the crystal lattice in this crystal form, we cannot entirely rule out that a longer ssDNA might occupy the dsDNA-interaction site B in the absence of a competing duplex DNA.

In SRAPd_3nt, the distance between the 3’ end of DNA-A and the 5’ end of DNA-B at the catalytic triad is around 3.2Å, which is sufficient to accommodate a phosphate group linking the two substrates together (**Fig. 2b**). Consistent with our observations, the affinity of SRAPd to dsDNA with a 3-nucleotide gap is ∼7-fold higher than intact dsDNA of the same sequence (**Supplementary Fig. 1**). These data suggest the potential for binding other types of gapped DNA structures that form during DNA repair (**Fig. 2d**), such as nucleotide excision repair intermediates.

Our SRAPd structures also shed light on other proposed activities of HMCES. First, proteomics studies using dsDNA baits with modified cytosines identified HMCES as a reader for oxidised 5-methyl-Cytosine (oxi-mC) containing duplex DNA^7^. The SRAPd only contacts one base-pair at the ssDNA-dsDNA junction (**Fig. 1b**); hence SRAPd of HMCES either recognizes a single oxi-mC at this junction or alternatively in single-strand regions. Second, HMCES was previously reported to recognize and cleave dsDNA with an oxi-mC modification in a metal ion dependent manner ^8^. Our structure and the conservation pattern do not reveal any capacity in the SRAPd for metal ion binding. Accordingly, we and others^4^ were unable to reproduce the HMCES nuclease activity. Taken together, our structures support an important role for HMCES in recognizing and sensing flapped and gapped DNA damaged products, and reveal its broad substrate recognition spectrum.

## ACKNOWLEDGMENTS

We are grateful to Dr. Haley Wyatt for fruitful discussions and comments about the manuscript. We thank Dr. Wolfram Tempel for collecting two datasets at the ALS beamline.

This research used resources of the Advanced Light Source, which is a DOE Office of Science User Facility under contract no. DE-AC02-05CH11231. Results shown in this report are derived from work performed at Argonne National Laboratory, Structural Biology Center (SBC) at the Advanced Photon Source. SBC-CAT is operated by UChicago Argonne, LLC, for the U.S. Department of Energy, Office of Biological and Environmental Research under contract DE-AC02-06CH11357.

## FUNDING

The Structural Genomics Consortium is a registered charity (no: 1097737) that receives funds from AbbVie; Bayer Pharma AG; Boehringer Ingelheim; Canada Foundation for Innovation; Eshelman Institute for Innovation; Genome Canada through Ontario Genomics Institute [OGI-055]; Innovative Medicines Initiative (EU/EFPIA) [ULTRA-DD: 115766]; Janssen, Merck & Co.; Novartis Pharma AG; Ontario Ministry of Research Innovation and Science (MRIS); Pfizer, São Paulo Research Foundation-FAPESP, Takeda and the Wellcome Trust. This research is also supported by the Canadian Institutes of Health Research [FDN154328 to CHA], intramural funds of the National Library of Medicine, NIH, USA to LA, and the National Cancer Institute (R35 CA210043 to AR).

## AUTHOR COTRIBUTIONS

L.H. performed the experiments, Y.L., M.R., and H.Z. cloned, expressed and purified the proteins. A.R., and C.H.A conceived the project. L.H., L.A., A.R., and C.H.A contributed to experimental design and review of data. L.H. and C.H.A wrote the paper.

## COMPETING FINANCIAL INTERESTS

The authors declare no competing financial interests.

## METHODS

### Protein expression and purification

Wild type and mutant variants of HMCES were subcloned into pNIC-CH vector by modifying the C-terminal tag with a TEV cleavable N-terminal His6-tag, and were expressed in *E. coli* Rosetta. All clones are sequence verified. The recombinant proteins were purified first by nickel-affinity chromatography and, after TEV cleavage of the His6-tag, by anion exchange and gel-filtration chromatography using S200 column. Purified SRAPd was concentrated to ∼20 mg/mL in 20 mM Tris-HCl [pH 8.0], 150 mM NaCl, 2 mM tris(2-carboxyethyl)phosphine (TCEP). The sequences for all cloned constructs were verified by sequencing, and the corresponding molecular weight for all purified constructs were verified by liquid chromatography-mass spectrometry LC-MS.

### Crystallization and structural determination

Apo-SRAPd was crystallized using sitting drop vapor-diffusion method by mixing 1:1 ratio of protein and reservoir solution containing 0.1 M Bis-Tris Propane, 2% Tacsimate, 20% PEG3350.

DNA used for co-crystallization were purchased from Integrated DNA Technologies, Inc. For co-crystallization, purified SRAPd protein at 12 mg mL^−1^ was mixed, at a molar ratio of 1:1.2, with different 3’ overhang DNA prepared by annealing equimolar amounts of two oligonucleotides, 5’-CCAGACGTT-3’ and 5’-GTCTTG-3’ for DNA_3nt; 5’-GTCTTG-3’ and 5’-CCAGACGTTGTT-3’ for DNA_6nt, and incubated for 0.5 h on ice. The mixture then crystallized using sitting drop vapor-diffusion method in a condition containing 25% PEG 3350, 0.2 M ammonium sulphate, 0.1 M Hepes pH 7.5 for SRAPd_3nt; and 20% PEG 3350, 0.1 M KCl, 0.1 M Bis-Tris pH 5.5, 0.05 M MgCl2 for SRAPd_6nt. Apo-SRAPd, SRAPd_3nt and SRAPd_7nt crystals were cryo-protected using reservoir solution supplemented with 20-30% glycerol and 20-30% ethylene-glycol, respectively, and cryo-cooled in liquid-nitrogen.

Diffraction data for the Apo-SRAPd, SRAPd_3nt and SRAPd_6nt were collected at the 19ID beamline of the Advanced Photon Source (APS), and 5.0.1 beamline of the Advanced Light Source (ALS) Berkeley Lab, respectively. Datasets were processed with XDS^11^ and merged with Aimless^12,13^. Initial phases for the Apo-SRAPd was obtained by molecular replacement with Phaser-MR^14^, using the (PDB ID: xxxx) as a search model. Initial phases for the two SRAPd-DNA complexes were obtained by molecular replacement with Phaser-MR^14^, using the Apo-SRAPd (PDB ID: 5KO9) as a search model. Models were built with COOT^15^, and refined with refmac5^16^. Data collection and refinement statistics are shown in (**Supplementary Table 1**). Figures were generated with PyMOL (http://pymol.org).

### Fluorescence-based DNA binding assay

All fluorescence polarization DNA binding assays were performed in a final volume of 20 µL in a buffer containing 20 mM Hepes, pH 7.4, 140 mM KCl, 5 mM NaCl, 0.1 mM Ethylenediaminetetraacetic acid (EDTA), 0.01% Triton X-100 and 0.2 mM TCEP in 384-well black polypropylene PCR plates. Fluorescence polarization (mP) measurements were performed at room temperature using a BioTek Synergy 4 (BioTek, Winooski, VT). The KD values were calculated by fitting the curves in GraphPad Prism 7.04 using nonlinear regression, one site-specific binding, equation Y=Bmax*X/(Kd + X). The sequences of 6-carboxyfluorescein (FAM)-labeled DNA oligonucleotides are listed in (**Supplementary Table 2**). All DNAs were purchased from Integrated DNA Technologies, Inc.

### Accession codes

Coordinates and structure factors have been deposited in the Protein Data Bank under accession codes: 5KO9 for Apo_SRAPd, 6NLD for SRAPd_3nt, and 6NLC for SRAPd_6nt.

