## Supplementary files for "Structural basis of HMCES interactions with DNA reveals multivalent substrate recognition"

Halabelian *et al.*

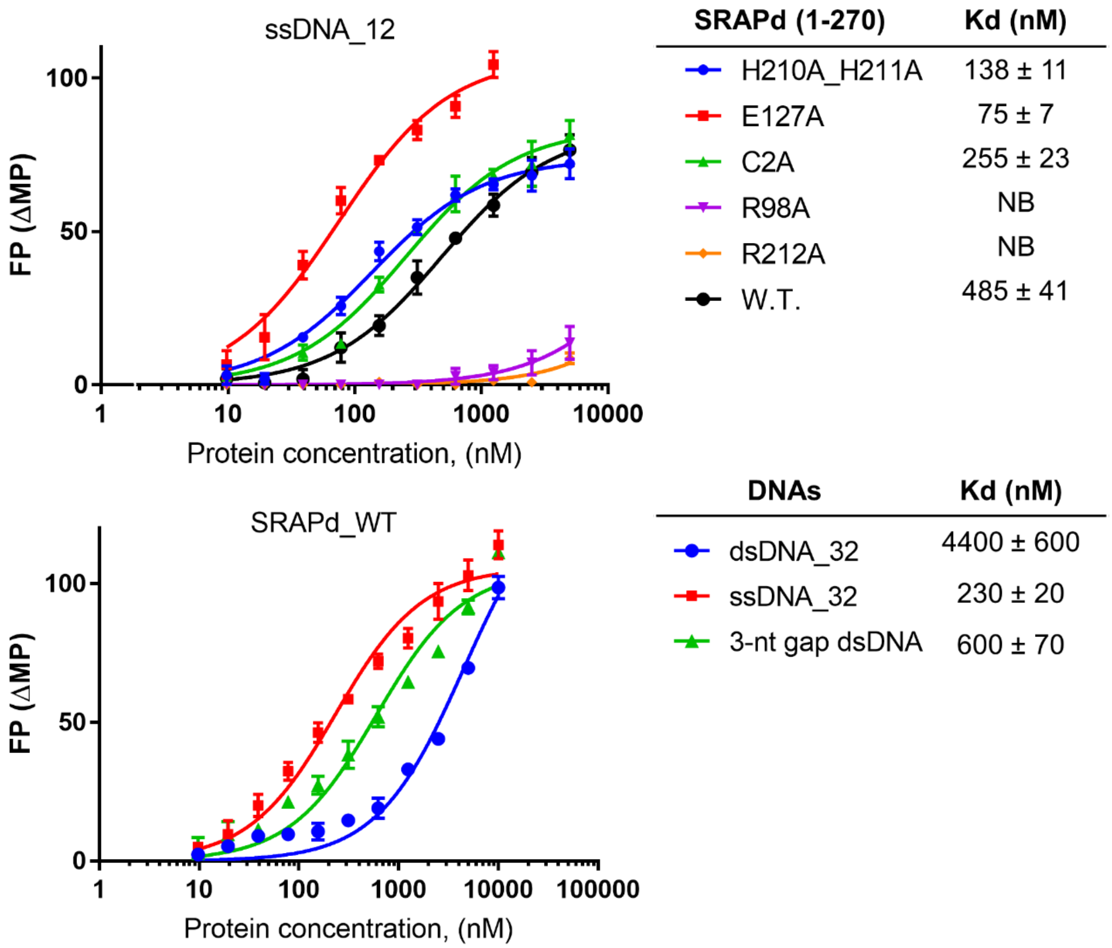

**Supplementary Figure 1. Fluorescence polarization DNA-binding assays for HMCES.** Comparison of ssDNA-binding activities of wild type (WT) and mutant SRAPd variants by fluorescence polarization (upper panel). Characterization of wild type SRAPd DNA-binding affinities to ssDNA, dsDNA, and dsDNA containing a three-nucleotide gap (3-nt gap dsDNA) (lower panel). NB, no detectable binding. Experiments were performed in triplicate.

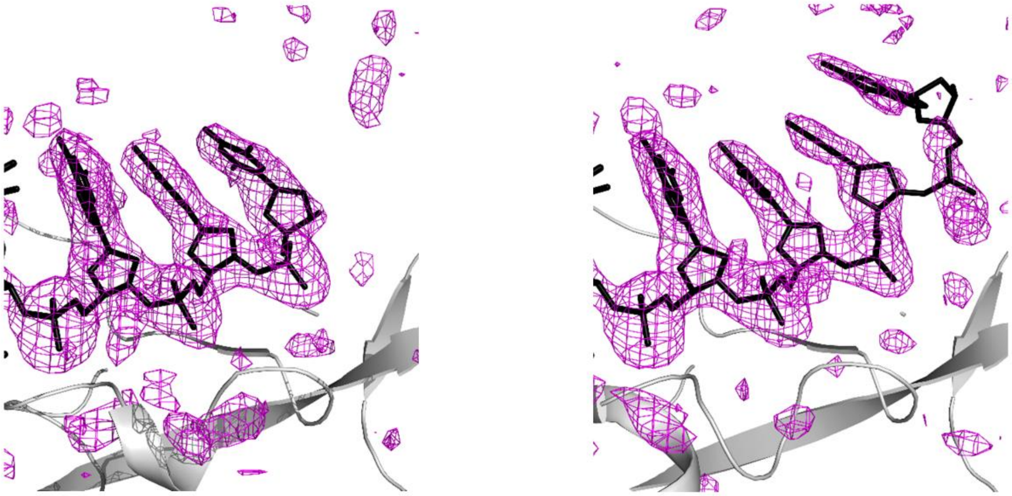

SRAPd_3nt SRAPd_6nt

**Supplementary Figure 2. Close-up view of the ssDNA binding cleft of SRAPd.** The $mF_o$-$DF_c$ electron density omit-maps for the ssDNA segment of 3' overhang DNA in SRAPd_3nt **(left panel)**, and SRAPd_6nt **(right panel)** are displayed as a magenta mesh, contoured at 2.5σ.

**Supplementary Table 1. Data collection and refinement statistics**

| | Apo-SRAPd (PDB ID:5KO9) | SRAPd_3nt (PDB ID: 6NLD) | SRAPd_6nt (PDB ID: 6NLC) |
|---|---|---|---|
| Data collection | | | |
| Space group | I 121 | I 121 | I 121 |
| Cell dimensions | | | |
| *a*, *b*, *c* (Å) | 80.10, 44.74, 82.90 | 55.73, 51.15, 149.21 | 55.86, 52.06, 148.31 |
| α, β, γ (°) | 90.00, 107.15, 90.00 | 90.00, 92.77, 90.00 | 90.00, 93.11, 90.00 |
| Resolution (Å) | 48.37-1.5 (1.53-1.50) | 48.38-2.05 (2.11-2.05) | 49.11-2.00 (2.05-2.00) |
| $R_{sym}$ or $R_{merge}$ | 0.045 (0.679) | 0.083 (1.046) | 0.063 (1.045) |
| $I / \sigma I$ | 15.9 (1.9) | 9.6 (1.2) | 11.3 (1.3) |
| Completeness (%) | 97 (93.2) | 99.8 (99.7) | 99.8 (99.8) |
| Redundancy | 3.8 (3.4) | 4.4 (4.4) | 4.6 (4.7) |
| Refinement | | | |
| Resolution (Å) | 48.37-1.5 | 48.43-2.05 | 38.09-2.0 |
| No. reflections | 41504 | 25225 | 27471 |
| $R_{work}$ / $R_{free}$ | 0.181/0.209 | 0.213/0.261 | 0.220/0.273 |
| No. atoms | 2292 | 2468 | 2453 |
| Protein | 2062 | 2365 | 2381 |
| Ligand/ion | 44 | 31 | 23 |
| Water | 186 | 72 | 49 |
| *B*-factors | 25.3 | 50.1 | 59.5 |
| Macromolecule | 24.6 | 50.1 | 59.6 |
| Ligand/ion | 35.7 | 57.6 | 61.3 |
| Water | 31.5 | 46.7 | 53.3 |
| R.m.s. deviations | | | |
| Bond lengths (Å) | 0.011 | 0.007 | 0.007 |
| Bond angles (°) | 1.499 | 1.438 | 1.471 |

*Values in parentheses are for highest-resolution shell.

**Supplementary Table 2.** DNA sequences used in DNA-binding assays.

| | DNA sequences | |
|---|---|---|
| ssDNA_12 | 6FAM-5’-CTGACGCGTATC-3’ | 6FAM: 6-carboxyfluorescein |
| ssDNA_32 | 6FAM-5’-TCTTCTGGTCCGGATGGTAGTTAAGTGTTGAG-3’ | 6FAM: 6-carboxyfluorescein |
| dsDNA_32 | 6FAM-5’-TCTTCTGGTCCGGATGGTAGTTAAGTGTTGAG-3’<br>5’-CTCAACACTTAACTACCATCCGGACCAGAAGA-3’ | 6FAM: 6-carboxyfluorescein |
| 3-nt gap dsDNA | 5’-TCTTCTGGTCCGGATGGTAGTTAAGTGTTGAC-3’-6FAM<br>5’-GGACCAGAAGA-3’<br>5’-GTCAACACTTAACTACCA-3’ | 6FAM: 6-carboxyfluorescein |
